# Background modeling, Quality Control and Normalization for GeoMx RNA data with *GeoDiff*

**DOI:** 10.1101/2022.05.26.493637

**Authors:** Lei Yang, Zhi Yang, Patrick Danaher, Stephanie Zimmerman, Tyler Hether, David Henderson, Joseph Beechem

**Author notes:** Correspondence NanoString Technologies, Seattle, US.

## Abstract

**Background:** NanoString’s GeoMx Digital Spatial Profiler (DSP) RNA assay can measure mRNA from hundreds of regions of customizable shape and size, yet it gives unique challenge in Quality Control(QC) and normalizating due to the omnipresent background noise incurred by the non-specific probe binding, which could not be addressed by conventional methods.

**Results and discussion:** Using Poisson Background model, Background Score Test, Negative Binomial threshold model and Poisson threshold model for normalization from the R package **GeoDiff**, we perform tasks including size factor estimation, QC and normalization on GoeMx RNA assay data. They are shown to outperform conventional methods like Limit of Quantification for QC as to consistency/false positive rate and 75% quantile normalization as to eliminating technical variability and recovering true signal.

**Conclusions:** We present a statistical model based workflow for QC and normalizing GeoMx RNA data using **GeoDiff**, justified by statistical theory and validated by real/simulated data.

## Background

Spatial gene expression platforms, which measure gene expression within tiny regions from across the span of a tissue sample, have opened a frontier in biology. These platforms harness idiosyncratic chemistries that introduce distinct technical effects into their data; however, spatial data is often analyzed with tools developed for bulk or single RNA-seq data, leading to bias and statistical inefficiency. Here we derive data analysis methods addressing the important technical effects of the GeoMx platform for spatial gene expression.

Given a tissue sample, the GeoMx RNA assay can measure mRNA from hundreds of regions of customizable shape and size. Its ability to assay flexibly-defined regions comes from a unique chemistry. Probes are constructed of two parts: an oligonucleotide that hybridizes to a target mRNA sequence in the tissue, and a second oligonucleotide barcode specifying the gene identity. These two halves are connected by a photocleavable linker. Probes are flowed across a tissue, binding their targets wherever they lie; then, precisely focused UV light cleaves the probes within user-defined regions and releases the barcodes for counting by short-read sequencing.

This chemistry leads to three complications for data analysis, each creating pitfalls for analyses designed for other platforms. First, probes stick at low rates to biological material other than their intended mRNA target, leading to background counts. Background impacts the expression of genes, especially genes with very low expression, introducing bias if ignored. Background also complicates the task of calling a gene as definitively detected. Second, in very small regions, many genes will have expression levels dropping to zero or background-level counts. These near-zero counts become statistically unstable after log-transformation, which has been commonly favored by gene expression analysts since the early days of microarrays. Third, the size of regions sampled in a single experiment can vary from one to thousands of cells, leading to a much wider range of technical effects in the raw data.

Normalization of GeoMx data has thus far required great care, often forcing a choice between multiple unsatisfactory methods. To facilitate future analyses, we have established data-generating models for GeoMx, and we have models to derive methods for the fundamental operations of GeoMx analysis. Most importantly, we define a normalization procedure that removes all major technical effects and allows for log-scale analyses. We also define a suitable model for the background and a test for whether genes are detected above background, as well as a model for size factor estimation. This suite of tools is implemented in the **GeoDiff** R package.

The background model is the starting point of the **GeoDiff** analysis. While there is some previous attempt to model negative control probes using Poisson based model for data from NanoString nCounter [1] and DSP [2] platforms, none of them allow parameter estimation for each negative probe and each sample as in **GeoDiff**. With full parameterization, model diagnostics can be carried out and outliers and batch effect can be identified, which provides more insight to the data than previously possible.

The Quality Control(QC) is a prerequisite before gene expression analysis. In QC for DSP, one commonly asked question is: “whether a feature/Region of Interest(ROI) has good signal”. One traditional way to answer this question is to define a per ROI Limit of Quantitation(LoQ) cutoff, to determine whether a single target is above the background per ROI by the cutoff, then to calculate the overall pass rate for each target or ROI as the metric for flagging them[3]. This approach is very heuristic. In this paper we formally clarify the meaning of “a target/ROI has good signal” in the context of GeoMx RNA dataset, and provide a statistical test for features and a metric for ROIs for signal QC purpose, with necessary justification and validation.

Normalization, the endeavor of removing technical variation from the gene expression data by finite step operations, is usually performed before other gene expression analysis[4]. This tradition dates back to early days of microarray gene expression analysis[5]. Among those, scaling method[6, 7, 8], i.e., dividing gene expression by a sample specific factor is commonly used on RNA-seq data for its simplicity. The output of normalization procedure is called “normalized expression”. Currently, the most used scaling normalization method for DSP RNA data is 75% quantile normalization(or Q3 normalization as 75% quantile is also called 3rd quartile)[3, 9]. Another popular scaling normalization method is from [7], which tried to find the best scaling factor using is trimmed mean of M values (TMM) compared to simply using quantiles. Log transformation with base 2 or 10 is often further applied on the normalized data to stabalize heavy tail distribution[10] of gene expression and facilitate interpretation using ratios. To avoid −∞ in log transformation, 0 is usually replaced by a small positive number. Although some non-scaling method exists, the interpretation is usually more tricky. For example, [11] takes the Pearson residuals from “regularized negative binomial regression” as the normalized expression.

This scaling normalization and log transformation approach has caused two major problems in GeoMx DSP data analysis. First, the relation between observed counts and real expression often is not(purely) scaling. Instead, genes are subject to influence of the background as well as the scaling(size) factor. The background has more of a “shift” effect on the data, which can not be corrected by scaling alone. Second, there is no base to the choice of that small positive number to replace 0, and it also causes artificial bimodality in the results. Here, we address both problems by reframing the normalization problem as a parameter estimation problem using appropriate saturated model[12] with regularization on the normalization parameters. Empirical Bayes(EB) procedure[6] could be implemented to select a data driven regularization term. We show that this method corrects for the background as well as fixes the artificial spikes. We also show the empirical bayes shrinkage increase precision of normalized expression by leverage information across genes.

## Results and discussion

### Overview

Our workflow begins by modeling the negative probes and performing corresponding model diagnostics. After that, we use the background model as a reference to perform statistical test for features and estimate signal size factor. Finally, we use information gathered from these steps to perform normalization.

We explain our approach using a dataset of GeoMx Human Whole Transcriptome Atlas (WTA) RNA assay on diabetic kidney disease (DKD) vs. healthy kidney tissue, a dataset of GeoMx Human Whole Transcriptome Atlas RNA assay on cell pellet array containing 11 human cell lines, and a dataset of GeoMx Cancer Transcriptome Atlas (CTA) RNA assay on an FFPE cell pellet array of mixed HEK293T and CCRF-CEM cell lines.

The WTA diabetic kidney dataset is used for NanoString Spatial Omics Hackathon, This dataset consists of three normal tissue samples and four samples with diabetic kidney disease. The details of this dataset are in [13].

For the WTA cell pallet array data, we profiled a formalin fixed paraffin embedded (FFPE) cell pellet array containing 11 human cell lines with human WTA. We placed a range of sizes of circular ROIs from 50-360 *µ*m diameter on each cell line, with 2-4 replicate ROIs per cell line and size[14].

In the CTA cell mixture data, HEK293T and CCRF-CEM (Acepix Biosciences, Inc.) were mixed in varying proportions, and aliquoted into a FFPE cell pellet array. Expression of 1414 genes in 700 *µ*m diameter circular regions from the cell pellets were measured with the GeoMx platform, and each gene has 3 probes[15].

For all datasets, ROIs were illuminated on the GeoMx Digital Spatial Profiler and tags were collected and sequenced as previously described. FASTQ files were processed using the Nanostring GeoMx NGS Pipeline v2.0 as previously described. Reads were deduplicated by UMI and deduplicated counts for each ROI and probe were used for analysis[16, 14]. The methods described in this paper all require raw count data without any preprocessing.

### Poisson Background model

The Poisson background model takes a form of

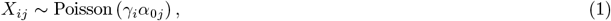

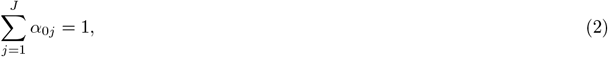

where *γ*_*i*_, *i* = 1, …, *I* are feature specific factors, and ***α***_0_ = (*α*_0*j*_), *j* = 1, …, *J* are sample size factors for each ROI. This model assumes, aside from random noise, variation in levels of *i*th feature in different ROIs is explained by the technical variation in the form of multiplicative size factor ***α***_0_. Therefore, this model is most suitable to features without biological variation and are subject to non-specific probe binding, i.e. negative control probes designed against synthetic sequences from the External RNA Controls Consortium(ERCC), which mimic the properties of mammalian sequences but have no homology to any known transcript. The constraint 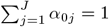 is imposed for identifiability. See Methods for further details.

Model diagnostics for Poisson Background model is available in **GeoDiff**. An empirical dispersion is calculated and a probability–probability(P-P) plot is generated in the diagnostics. The expected dispersion for Poisson distribution is 1. It is called overdispersion when the empirical dispersion is larger than 1, which is the most direct consequence of deviation from model assumption, likely caused by outliers or batch effect. The recommended remedy for outliers in negative probes is setting them to be missing and refiting the model. For batch effect we recommend people to find the root cause, as it impacts all aspects of data analysis. Fitting a Poisson Background model with a grouping variable is a workaround when batch effect is present(see Methods for details).

We fit the Poisson Background model without and with grouping variable indicating slide ID to the negative probes of WTA kidney dataset. The estimated dispersion are close to 1, and probability–probability(P-P) plot(Figure 1A, 1C) align with the diagonal line, showing both models are good fit for the data. This is further confirmed by the consistency of the feature factors across slides(Figure 1B, 1D).

**Figure 1:**
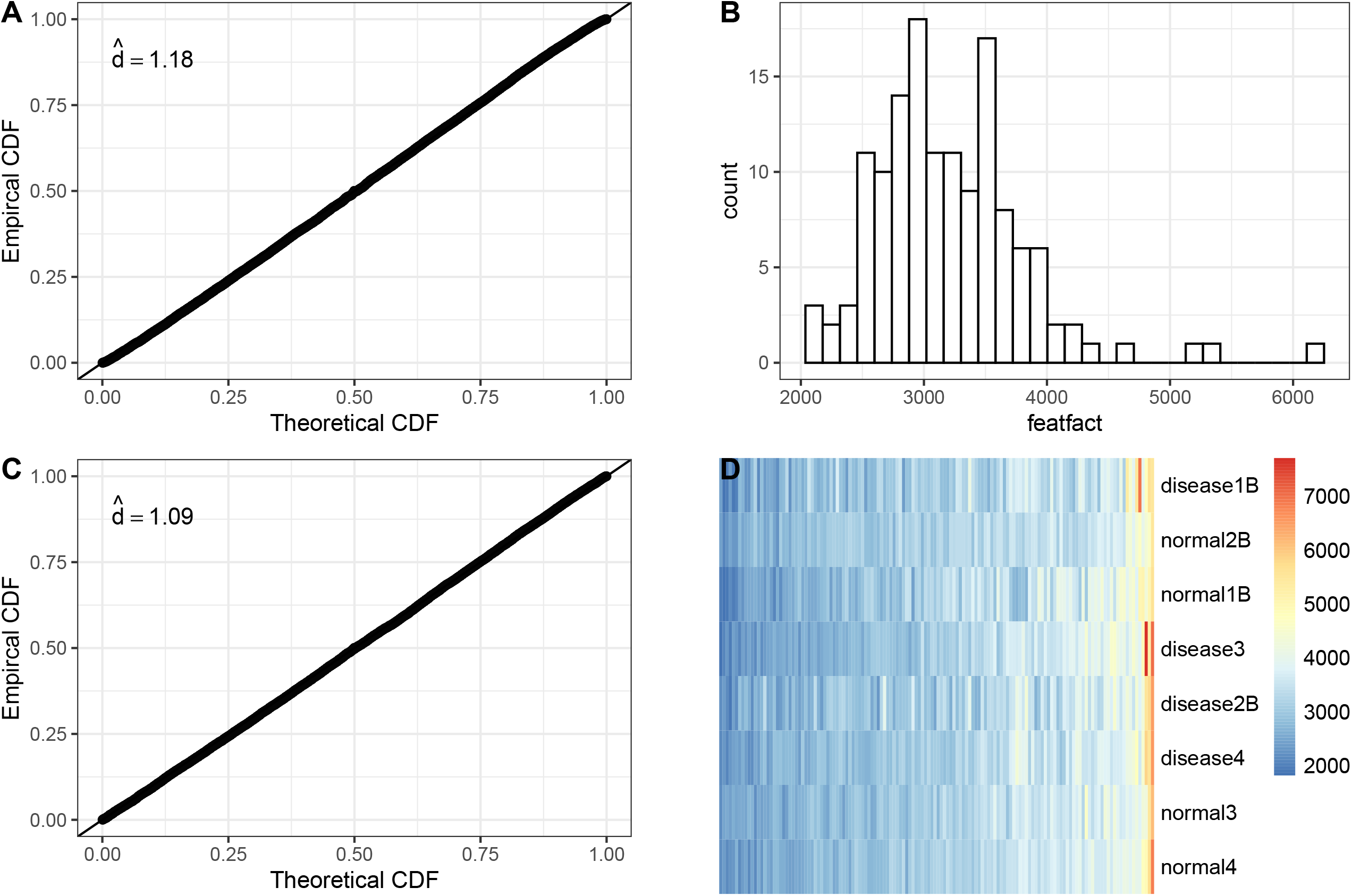
Diagnostics and distribution of feature factors from Poisson Background Model. Poisson Background model, treating the feature factors constant in the whole dataset(A, B), or within each slide(C, D). The P-P plot (A, C) show both models have a good fit with estimated dispersion close to 1. This can be explained by homogeneous distribution of the slide specific feature factors (D).

### Background Score Test

After applying the Poisson Background Model to the negative probes, it is imperative to test whether a target is above the background using the fitted model as a reference. It is called Background Score test for that it is derived using the score test paradigm(also known as the Lagrange multiplier test in econometrics[17]).

There are two versions of Background Score Test derived without and with a (Gamma distribution) prior assumption for feature factors.

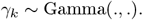

Background Score Test with prior takes into account the variation between feature factors thus is recommended and set as default. The “with prior” version is also implemented for all Background Score Test in this manuscript. For details of Background Score Test without and with prior, refer to Methods for details.

We compare the Background Score Test to the current default feature QC method LoQ, defined as the geometric mean times exponential of two standard deviations of the logarithm negative probes[3], in both the case study and simulation study.

#### 0.0.1 Case study

In this analysis, we apply both Background Score test and LoQ method to the WTA cell pallet array data described above by five different ROI sizes, including 50, 80, 110, 250, 360 *µ*m. The Background Score test with prior is applied to raw count data to extract the p values. In contrast, we flag the gene above the background if it exceeds LOQ in at least one ROI. The heatmap displays how −log(p value) from background score test and indicators on whether above background (1: Yes; 0:No) from LOQ change over increasing ROI sizes (Figure 4).

**Figure 2:**
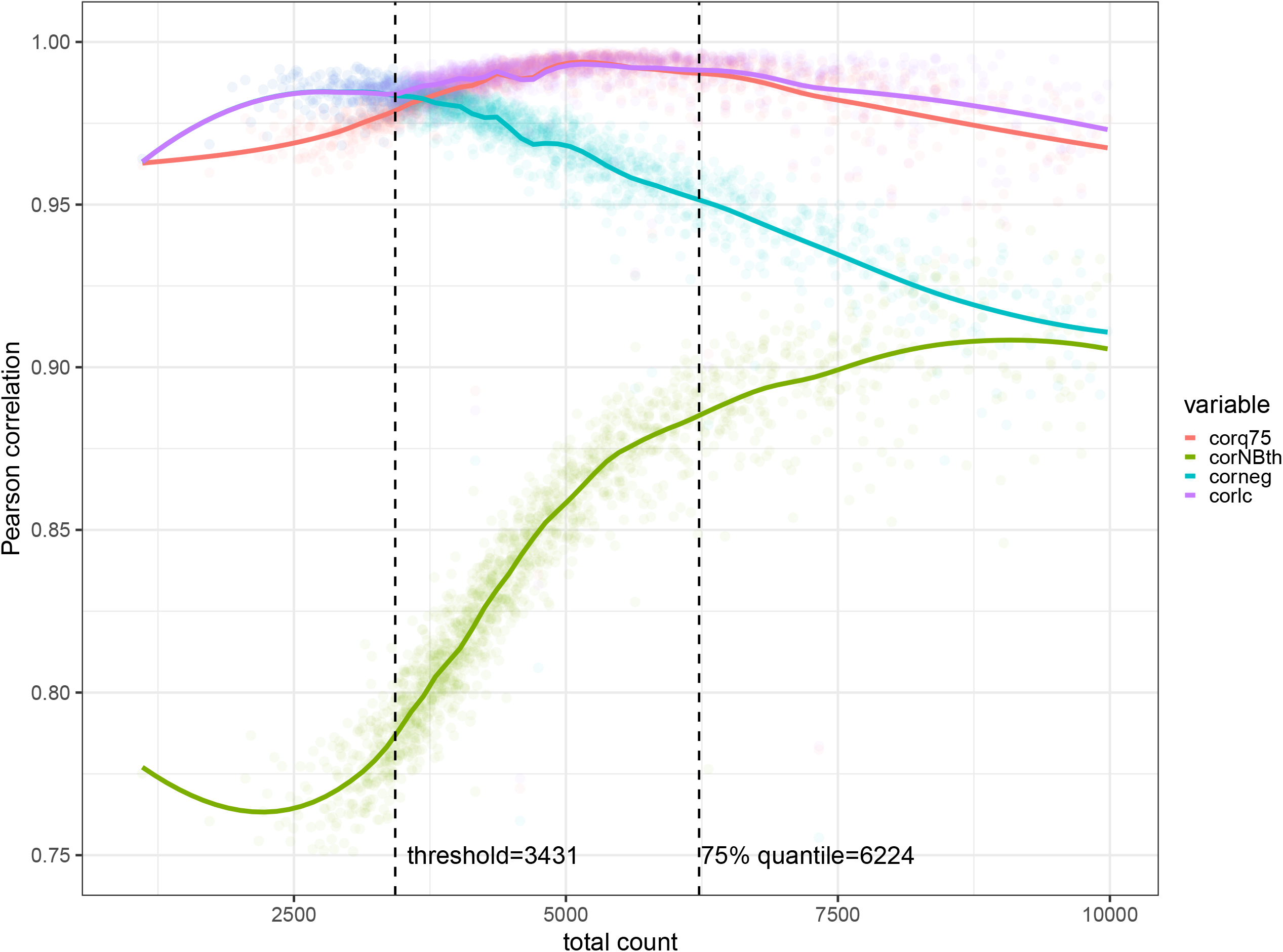
Correlation of counts and size factors. This plot shows the correlation of size factors: background sizefactor estimated from Poisson Background Model(corneg), signal sizefactor estimated from Negative Binomial threshold model(corNBth), linear combination of both based on the Negative Binomial threshold model(corlc), and 75% quantile(corq75) with average counts of binned features(bin size=10) sorted by scores of their Background Score Test.

**Figure 3:**
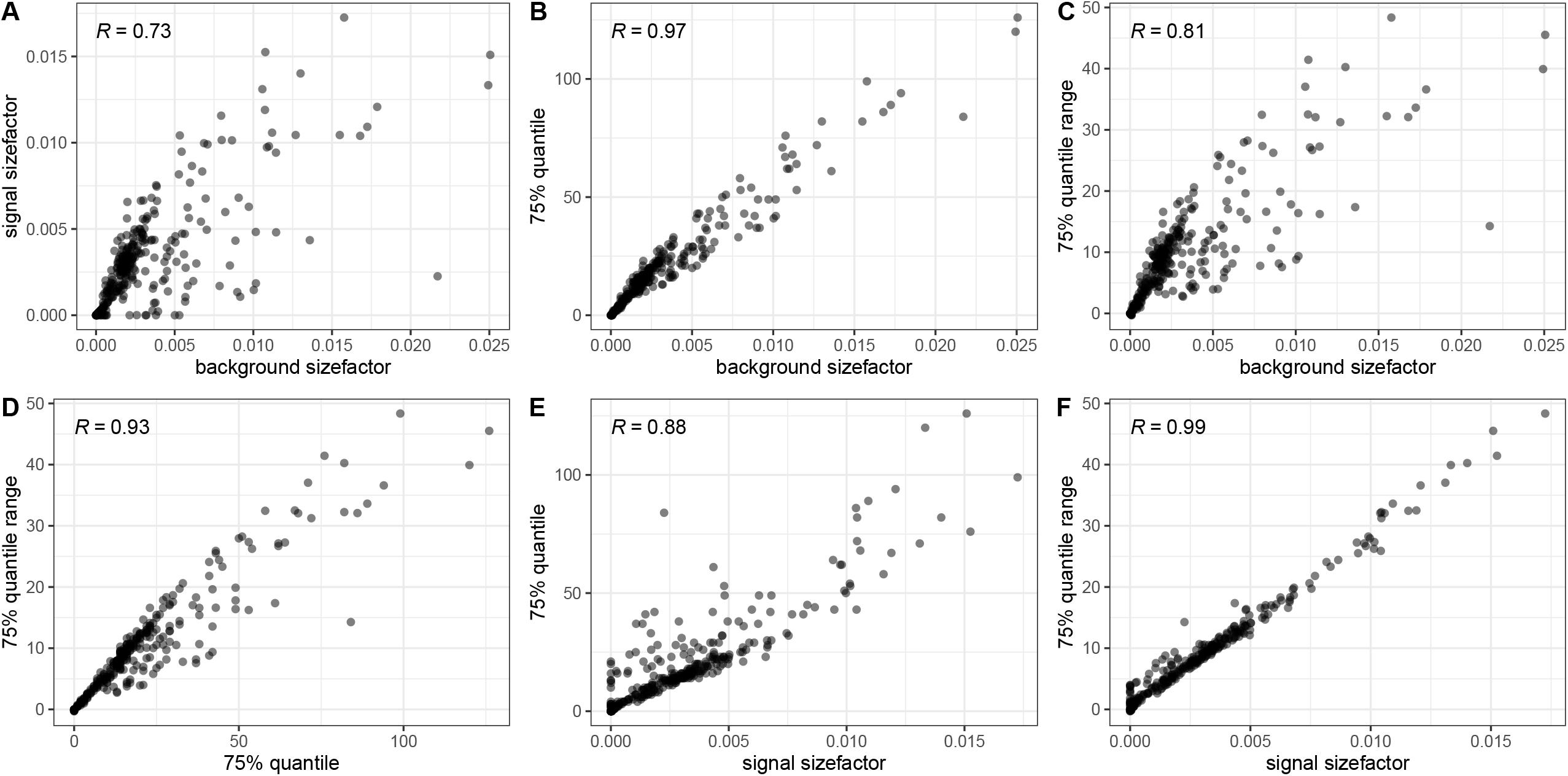
Scatterplots between size factors, 75%quantile and 75% quantile range. Scatterplot between background size factor vs signal size factor(A), 75% quantile(B) and 75% quantile range(C); between 75% quantile(B) and 75% quantile range(D), and signal size factor vs 75% quantile(E) and 75% quantile range(F).

**Figure 4:**
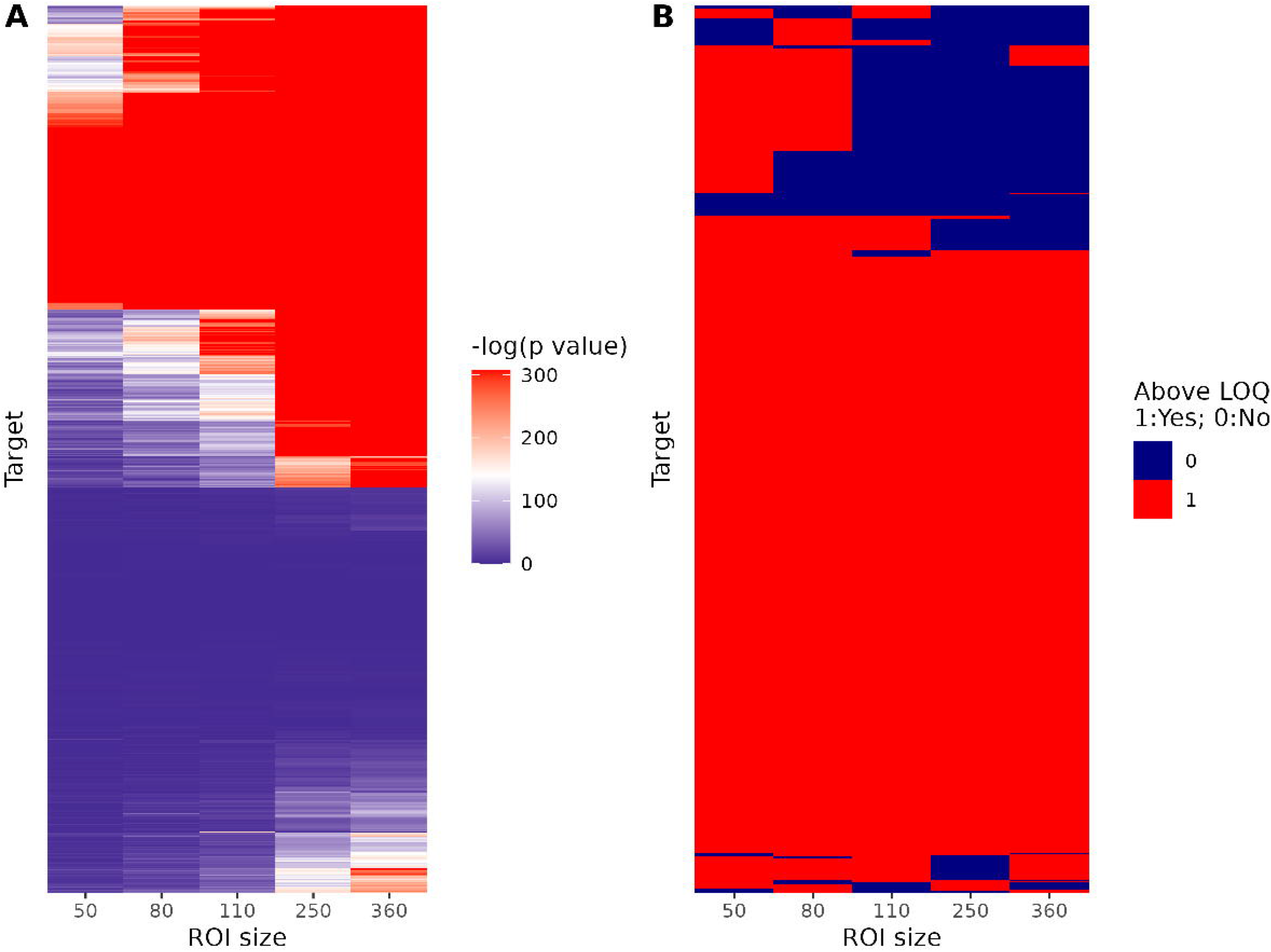
Heatmaps of p-value and indicators of targets above Limit of Quantitation (LOQ) across different ROI size in WTA cell pellet array data. A:Each row of the heatmap represents a target’s −log(p-value) from background score test across different ROI sizes. B:Each row of the heatmap represents the indicator of whether a target is above LOQ (1: above; 0: below) across different ROI sizes.

It is notable that Background Score test yield p values consistently increase with ROI sizes, meaning that it is more likely to be detected above background in larger ROI size. In contrast for LoQ, some genes are present in smaller ROI sizes but not larger ROI sizes, shows the LoQ method is less stable and more prone to be influenced by random noise.

Transcripts per million (TPM) from RNA-seq for different cell lines was used as a reference. We have shown that score statistics have better correlation with TPM values, measured in Spearman correlation, compared to the proportion of ROIs with gene expression above the LOQ (Figure 5).

**Figure 5:**
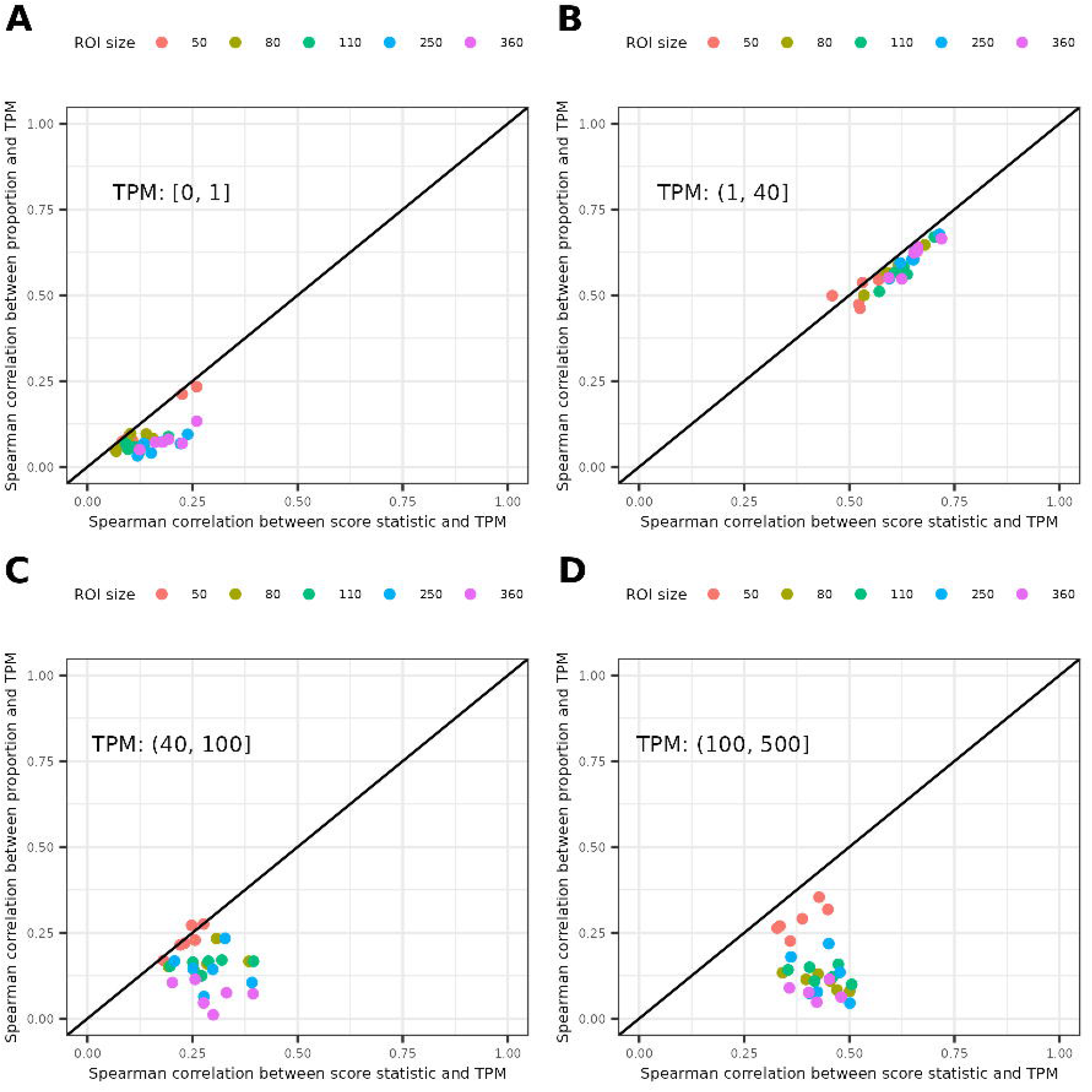
A comparison of Spearman’s correlations between Transcripts Per million (TPM) and score statistics vs. TPM and the proportions above LOQ. Comparison is split at varying TPM intervals, including (A) [0, 1], (B) (1, 40], (C) (40, 100], and (D) (100, 500], in WTA cell pellet array data.

#### 0.0.2 Simulation study

The simulation study evaluates the false positive rates (FPRs) based on the 136 negative probes from the WTA cell pallet array data. For each ROI size, we estimate the size factors and feature factors and randomly sample 10,000 negative probes for each ROI. Then, the raw counts of genes are drawn from a Poisson distribution with mean of the size factor multiplied by the feature factor. Similar to the case study, we applied both Background Score test and LOQ method on the simulated population of negative probes (n = 10,000). False positive gene is defined as a gene where the null hypothesis is falsely rejected and the proportion of the total false rejected genes as the FPR. By setting the nominal level to be 0.001, we observe that LOQ method yields much higher FPRs in smaller sized ROIs, i.e., 50 *µ*m, 80 *µ*m, and 110 *µ*m. It is consistent with the finding where LOQ only detects a subset of genes in smaller-sized ROIs (Figure 6), which could be false positives.

**Figure 6:**
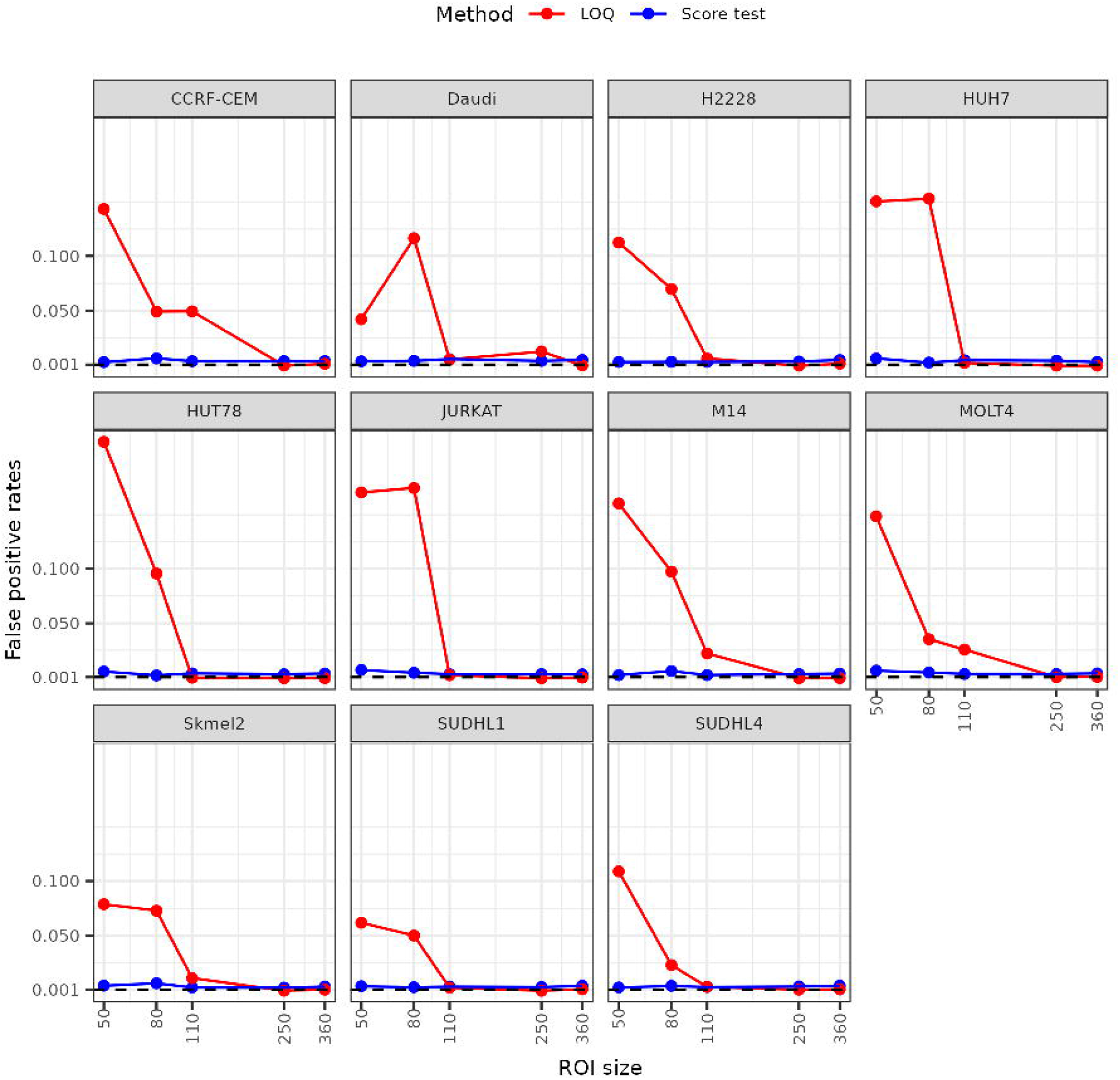
False positive rates (FPRs) of Background Score test and LOQ method for varying ROI sizes and different cell lines in simulated negative probes. The two solid lines represents the FPRs of background score test (blue) and LOQ method (blue), respectively. The dashed line represents the threshold of nominal alpha level, 0.001.

### Negative Binomial threshold model: The signal size factor

In GeoMx RNA data, the size factor of targets are usually not the same as the size factor of the background, as is demonstrated in Figure 2, the correlation of features with background size factor decreases as the abundance increases. To estimate the signal size factor with background taken into consideration, we fit the following Negative Binomial threshold model to a set of “high genes/features” or “above the background genes/features”, defined as features above the background determined by the Background Score Test, and assuming the real expression level of these gene is constant across all ROIs.

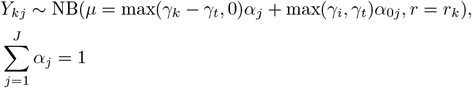

We use Negative Binomial distribution due to the overdispersion brought by the biology heteogenity as well as the “oversimplication” of the constant assumption, which is necessary for deriving size factors and is widely used for scaling normalization method[6, 7, 8]. For example, the commonly used 75% quantile normalization is implicitly assuming genes are roughly constant after dividing by the 75%. In [6], a median-of-ratios method is used to derive the size factor, which implicitly assumes the ratios are roughly constant for all genes. Here we explictly make this assumption using a representative set of features above the background.

The model is fitted by taking the estimated background size factor 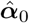 as input. The threshold *γ*_*t*_ can be either set to be the mean of background feature factor 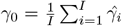 or unspecified and estimated from the model. Since *γ*_*t*_ ≈ *γ*_0_, when *γ*_*t*_ is estimated, it should be within 20% of *γ*_0_, otherwise the model might not be a good fit for the particular dataset. Other parameters *γ*_*k*_, *r*_*k*_, ***α*** are also estimated from the model.

An important implication of this model is that the size factor of each feature is defined as a linear combination of background size factor and signal size factor depending on its abundance as

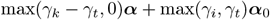

For low abundance features, it basically behaves the same as the background size factor; as the abundance gets higher, the linear combination is weighted heavier towards the signal size factor.

By the constraint 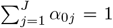 and 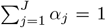, the *γ*_*k*_ is approximately the total count of *k*th feature, since if we use Poisson distribution instead of Negative Binomial distribution in the model, the total count of *k*th feature is the maximum likelihood estimator of *γ*_*k*_. With this knowledge, the linear combination is easy to calculate for each feature without applying the model on all of them.

Figure 2 shows this linear combination has the best correlation with features from different abundance ranges among all size actors to be compared, including background/signal size factor alone and 75% quantile. The performance of 75% quantile is comparable for features with moderate abundance, but is considerably worse for low/high adundance genes. The background sizefactor has the same correlation with low abundance features as the linear combination, but the correlation deteriorate as the abundance increases; while the signal sizefactor has the overall worst correlation but the correlation increases as the abundance increases.

### 0.1 ROI signal metric

The signal size factor ***α*** is an estimator for ROI signal levels, which can be naturally used to flag ROI with low signal. However this should not done directly on ***α*** since they are scaled to sum up to 1, so on average the more samples the lower the value of each element of ***α***. We could define signal level of ROI by

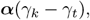

for *γ*_*k*_ representing a certain quantile of genes. One can just use 75% quantile. To encourage keeping ROIs, one can also use a higher quantile, such as 90% quantile. Then this metric represents the expected above-the-background counts of certain quantile of features.

However, ***α*** is defined and estimated via a model, which is not intuitive and straightforward for some users. We propose using the following metric named “quantile range” as a proxy. Denote 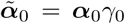 (see Methods for rationale), the 75% quantile range is defined as

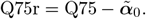

Q75r is just the difference between 75% quantile and the average background level. Quantile ranges are shown to be better correlated with 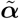 than quantiles (Figure 3) so that it can be used as an substitute for ROI signal metric for QC purpose without fitting the model.

### Poisson threshold Normalization

From a statistical perspective, normalization is to fit a saturated model in which there is one parameter(log_2_ normalized expression) for each observation. This is called saturated since the number of parameters is identical or larger than the number of obervations[12].

Assume ***β***_*k*_ = (*β*_*k*1_, …, *β*_*kJ*_) be the vector of log_2_ real expression of *k*th feature for *J* samples. The goal of normalization is to optimally estimate ***β***_*k*_ with all available information. There is no overdispersion caused by lack of information in saturated model, so Poisson distribution is applied.

For traditional scaling-normalization, we fit the following Poisson model

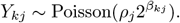

With observed *Y*_*kj*_ and any size factor ***ρ***, the normalized expression is equivalent to

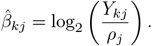

Following the same argument for Negative Binomial threshold model, a Poisson threshold model is more appropriate for GeoMx RNA data. The size factors calculated by Poisson Background model and Negative Binomial threshold model need to be rescaled to stay invariant to constraints (see Methods for more details).

Let 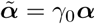(or *γ*_t_***α***_0_) and 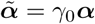

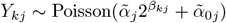

One direct consequence of such model is the observations will be fitted perfectly without constraint or priors, and when *Y*_*kj*_ = 0, the MLE of *β*_*kj*_ is −∞. But perfect fitting is usually suboptimal due to overfitting. Parameter regulation is desired for getting parameters generalize better as well as restrict infinite parameter estimation, so we specify a prior for the in addition to the model

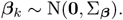

The precision matrix 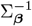 is estimated by a 2-step Empirical Bayes procedure: in Step 1, the model is applied to high abundance features with a default weak prior; in Step 2, a precision matrix is calculated using the estimated parameters, then the model is applied to all features of interest, using the estimated precision matrix.

There are two options for calculating the precision matrix in **GeoDiff**. One option is “equal”, which simply calculates the empirical precision matrix of estimated parameters; one option is “contrast”, which simply calculates the precision matrix with only contrast information from the estimated parameters. Precision matrix estimated by the “contrast” option avoids the abundance information of selected high abundance features influence model fitting other features by using the contrast information only, thus is recommended and set as default in **GeoDiff**. Both “contrast” and “equal” way of estimating precision matrix is parameterization invariant (see Methods for details).

The 2-step Empirical Bayes procedure can be applied for whole dataset or separately for each level of a grouping variable. The usual choice of grouping variable is the slide ID. When fitted without grouping variable, ROIs in different slides could influence each other’s normalized expression, thus the option of a grouping variable is recommended for normalization in multiple slides data.

#### 0.1.1 Case study: WTA diabetic kidney dataset

We evaluate the performance of this approach to other normalization methods by WTA diabetic kidney dataset and CTA cell mixture dataset.

For WTA diabetic kidney dataset, we apply different normalization methods and the distribution of their log_2_ normalized expression in slide “disease4” are compared in Figure 7. The Poisson threshold normalization(Figure 7E, Figure 7F for using slide ID as grouping variable) can align gene expression of different ROI much better than other methods. Specifically, the scaling normalization methods (Figure 7B, Figure 7C) do not correct for shift and the wiggles on the lower end while the scaling method with background subtraction (Figure 7D) incur a big spike on the lower end due the negative numbers it induces, both Poisson threshold normalization (Figure 7E, Figure 7F) can align the distribution of gene expression in a unimodal fashion.

**Figure 7:**
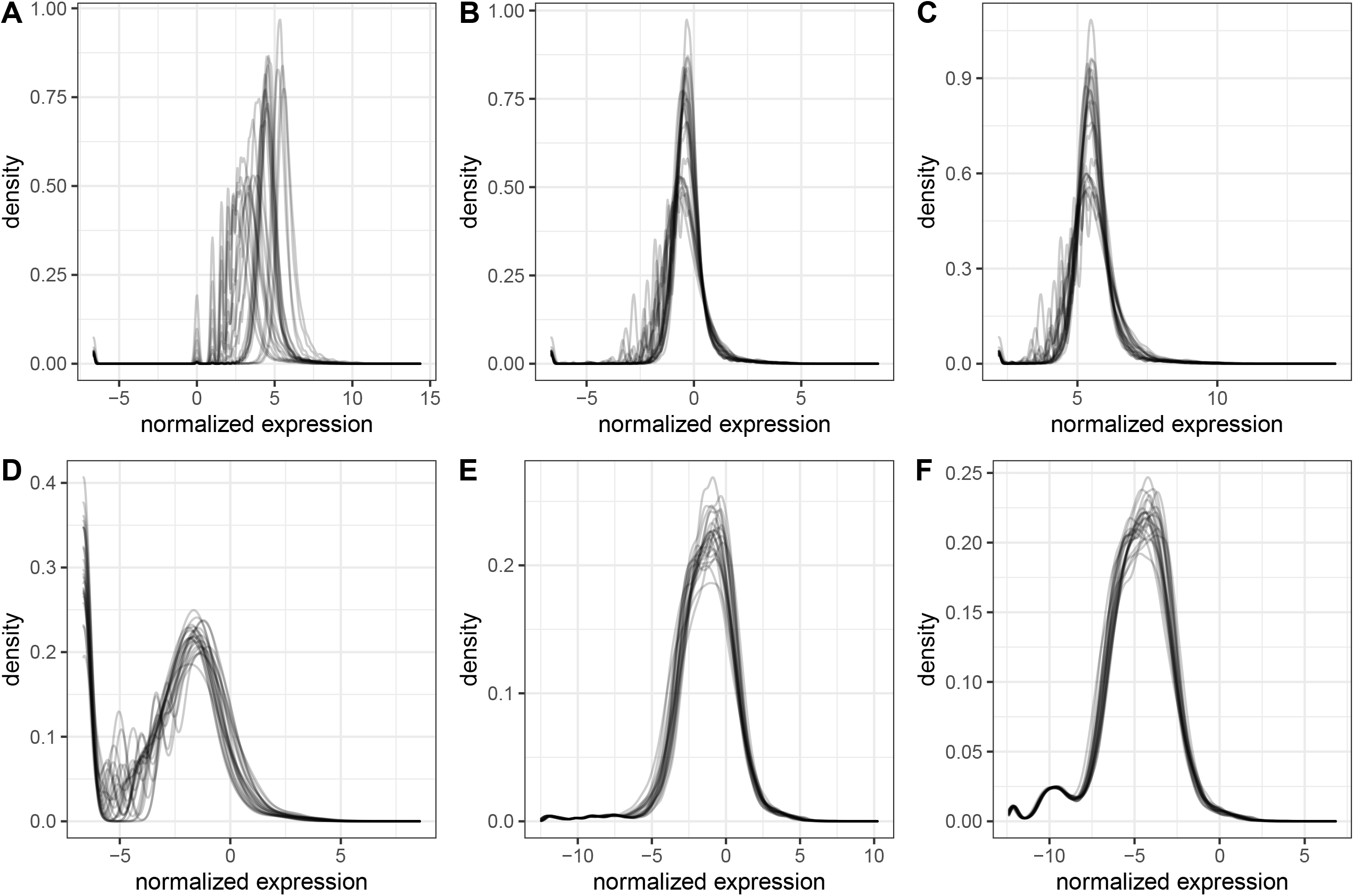
Density plot for gene expression for slide “disease4” in kidney data. the density of log2 expression of raw counts(A), 75% quantile normaliztion(B), TMM[7](C), 75% quantile normaliztion with background subtraction(D), Poisson threshold normalization(E) and Poisson threshold normalization by grouping variable (F) for the slide “disease4”.

Furthermore, PCA plot Figure 8 shows Poisson threshold normalization better removes technical variability and reveals better clustering pattern in terms of grouping variable and biological variable than 75% quantile normalization for “middle” abundance genes, defined as genes between the 40% and 60% quantiles of scores from the Background Score Test. “middle” abundance genes are above the background as indicated by the Background Score Test with *p* < 1*e* − 3, but they are still heavily influenced by the background. 75% quantile normalization yield results with its PC1 dominated by technical variation defined as the log_2_ ratio of 75% quantile and background size factor, showing it is not able to remove this technical variability, while PCs in both Poisson threshold normalization are less dictated by this variable, showing they both remove this technical variability better. Poisson threshold normalization results both cluster better to the grouping variable and the main biological variable: region. Interestingly, the Poisson threshold model without grouping variable yield better overall clustering in terms of ROI regions with apparent and not distinct clustering in terms of slides, while the Poisson threshold model with slide ID as grouping variable yield better overall clustering in terms of slides, with ROI regions clustered in the same direction within the same slide.

**Figure 8:**
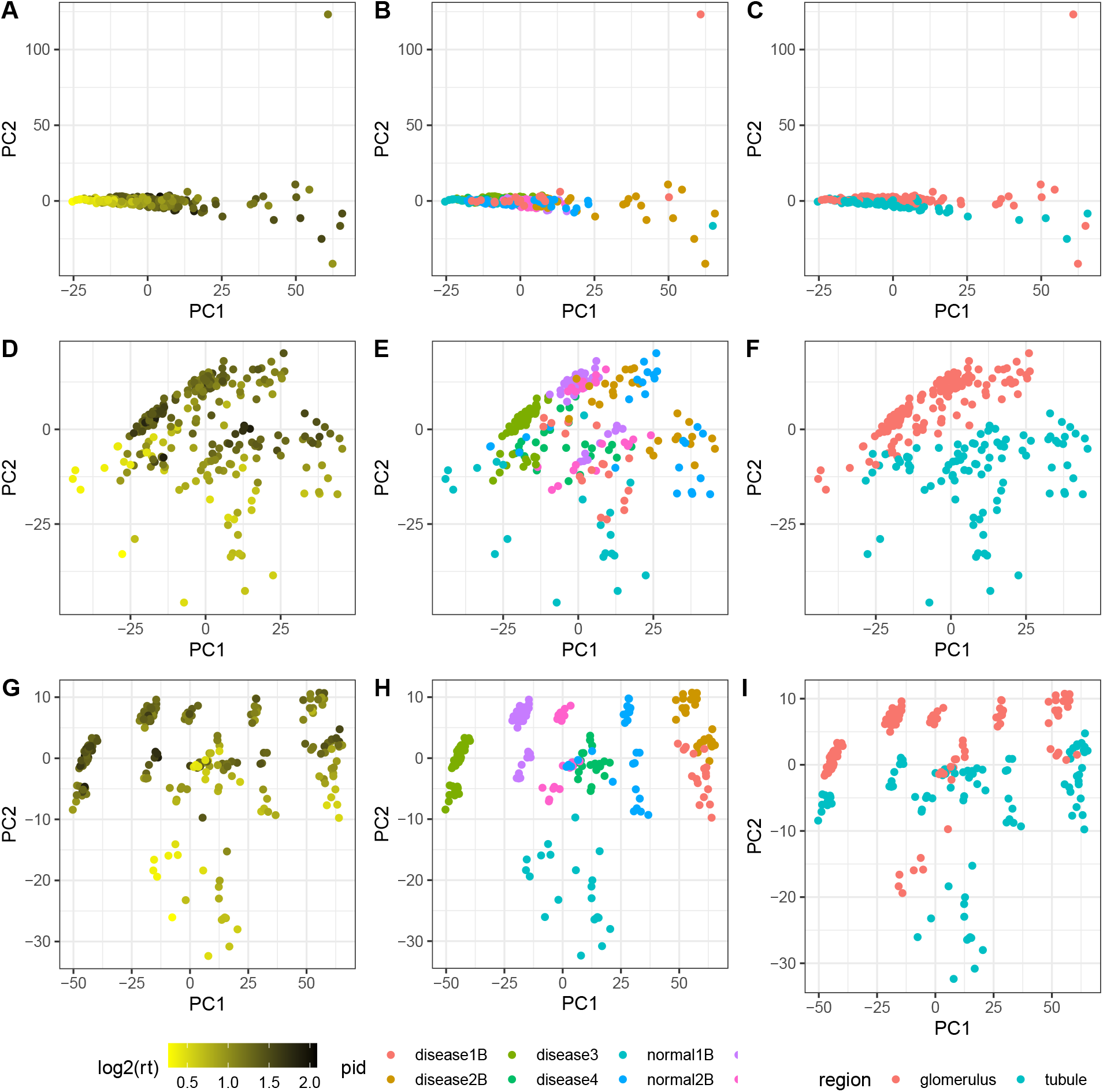
PCA plot for normalized expression of middle abundance genes. Figures of first two principle components of normalized expression of 75% quantile normalization(A, B, C), Poisson threshold normalization(D, E, F) and Poisson threshold normalization by grouping variable(G, H, I) on “middle” abundance genes, colored by the log_2_ ratio of 75% quantile and 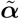 representing a key technical variation (A, D, G), slides ID (B, E, H) and ROI regions (C, F, I).

#### 0.1.2 Case study: CTA cell mixture dataset

For CTA dataset of mixture of HEK293T and CCRF-CEM cell lines with varying proportions, we has “partial truth” we can leverage. Specifically, while we do not know the “true log_2_ expression” of every gene of HEK293T and CCRF-CEM, if we assume *τ*_*k*1_, *τ*_*k*2_ are the “true log_2_ expression” of *k*th gene of HEK293T and CCRF-CEM, then the “true log_2_ expression” of any mixture with the proportion of HEK293T as *x* is

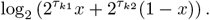

By the following steps, we evaluate how normalized expression from different methods align with this unique functional relationship.

1. Perform normalization
2. For *k*th gene, estimate *τ*_*k*1_, *τ*_*k*2_ using the log_2_ normalized expression ***β***_*k*_ by minimizing mean squared error (MSE)

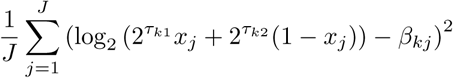
3. For *k*th gene and *j*th ROI, where *x*_*j*_ is the proportion of HEK293T, calculate the expected log_2_ normalized expression by

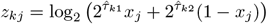
4. Calculate expected raw count by the model assumption of Poisson threshold normalization and 75% quantile normalization by the Poisson threshold model and Poisson scaling model assumptions.
5. Calculate the correlation of expected raw count vs the raw count

Figure 9A showcases this unique functional relationship between true log_2_ expression and percentage of HEK293T in the mixture using gene ABCB1 as example, in which 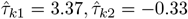 estimated by Poisson threshold normalization results. Figure 9B shows how 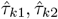 are fitted by the steps 1,2 and Poisson threshold normalization with and without EB prior gives very close estimates but the MSE is smaller in Poisson normalization with EB prior, which is further confirmed by Figure 9C where MSE of all genes from the two models are compared. Thus we can say the EB prior of Poisson threshold normalization imposed information implied by high features to all features and yield more precise results.

**Figure 9:**
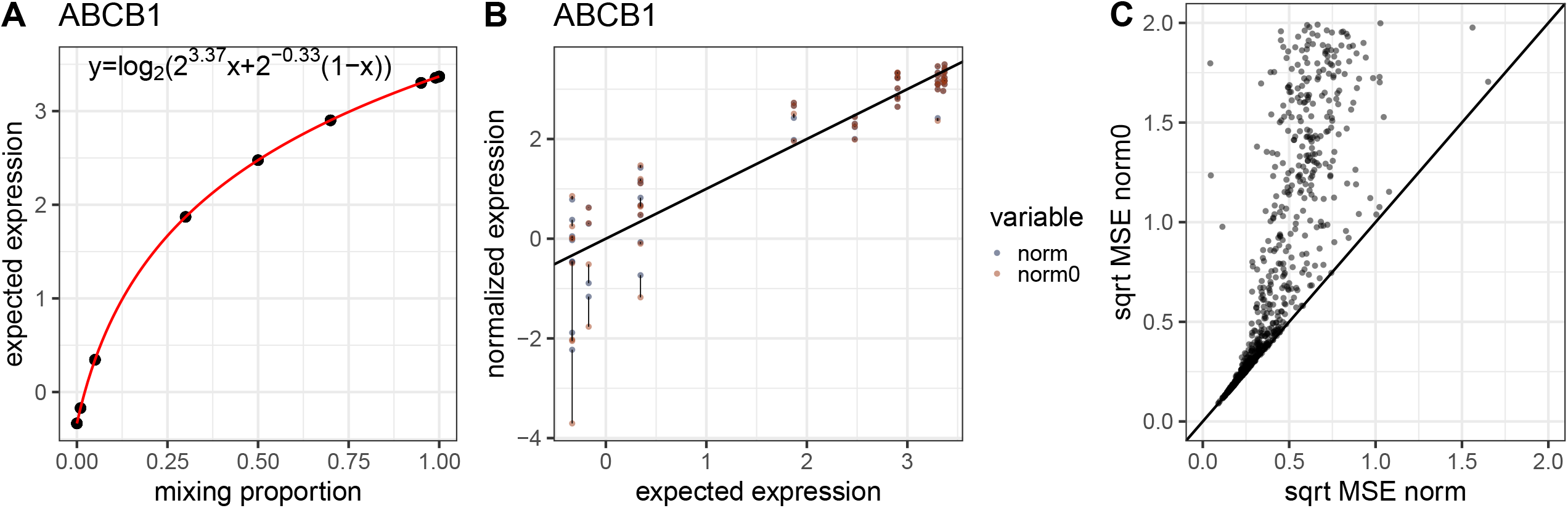
The relation between true log_2_ expression and HEK293T percentage. Using gene ABCB1 as an example(A, B), taking its estimated log_2_ expression of HEK293T and CCRF-CEM as true, the expected log_2_ expression for any mixture of the two with the percentage of HEK293T *x* follows a functional form as in A, and B shows the distance of log_2_ expression from Poisson threshold normalization without and with EB prior and the calculated expected log_2_ expression, in which Poisson threshold normalization with EB prior gives results closer to the expected values for ABCB1; C shows for all genes, the square root of MSE of estimating the log_2_ expression of HEK293T and CCRF-CEM using the Poisson threshold normalization without and with EB prior.

The results of Step 4 of the workflow is shown in Figure 10, in which correlation of expected raw count from different normalization methods vs raw count are compared. In Figure 10A, Poisson threshold normalization with and withour EB prior are compared; Figure 10B, Poisson threshold normalization with EB prior and 75% quantile normalization are compared, and Poisson threshold normalization with EB prior has better performance in both comparison.

**Figure 10:**
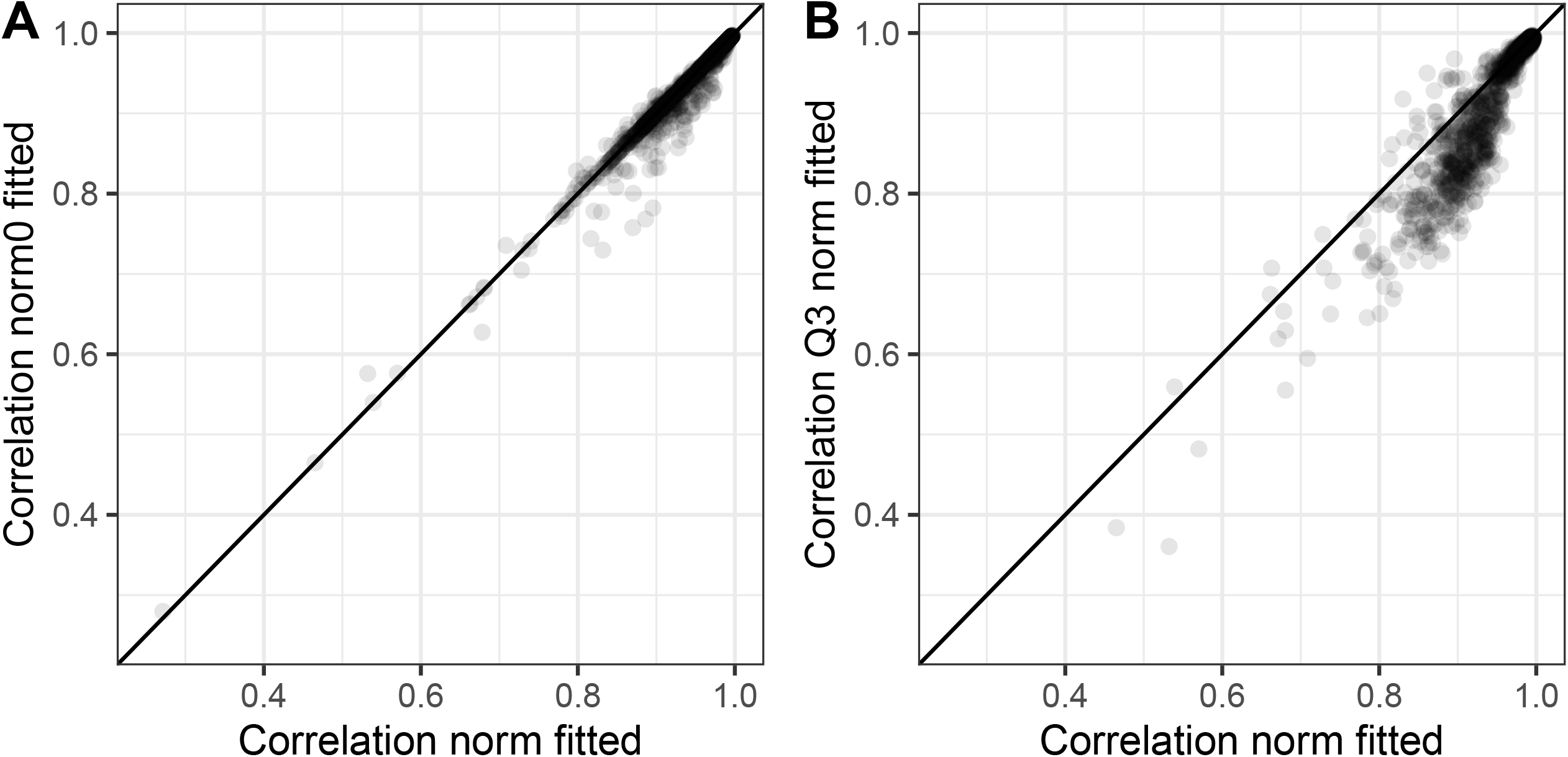
Correlation of expected expression vs the raw counts. A: the correlation of expected raw count from Poisson threshold normalization vs raw count, the X axis is with EB prior and the Y axis is without EB prior. B, the correlation of expected raw count from Poisson threshold normalization with EB prior(X axis) vs raw count and 75% quantile(Y axis) vs raw count.

### Targets with multiple probes

There are products on GeoMx platforms with multiple probes for one target. When the number of probes are big, some screening can be conducted to ensure the probes correlate well with each other and are in similar dynamic range. After that, the sum of multiple probes for that target is used for analysis, as Poisson and Negative Binomial distributions have the following property.

For

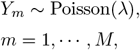

then

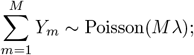

and

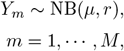

then

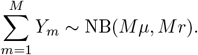

Furthermore, the sum of counts are sufficient statistics[18] for both models so no information is lost in the aggregation counts for parameter estimation. As long as the number of probe involved in the probe sum is correctly specified in the models, the estimated parameter will be scaled properly. The CTA cell line mixture dataset has 3 probes for each target and has been aggregated as described. The previous section shows the **GeoDiff** methods work well in such case.

## Conclusions

This paper describes statistical models/tests from **GeoDiff** including Poisson Background model, Background Score Test, Negative Binomial threshold model and Poisson threshold model for normalization, and recommends a GeoMx RNA data analysis workflow starting with modeling the negative probes for background size factor, followed by modeling the high features for estimating signal size factor, testing whether a target is above the background, assessing the signal level of an ROI, and perform normalization using outputs from these models/tests. We have shown the proposed methods outperform conventional methods in various aspects.

## Methods

### 0.2 Identifiability

The Poisson Background model is intrinsic non-identifiable, i.e. different sets of parameters *cγ*_*i*_, *α*_0*j*_*/c* for any positive constant *c* yield the same model. To impose identifiability, a constraint like 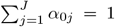 must be enfored. Constraints are arbitrary. However, 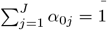 is equivalent to 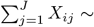 Poisson (*γ*_*i*_), making the total count 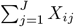 the MLE of *γ*_*i*_. This interpretation is convenient in a lot of applications.

The Negative Binomial threshold model suffers from identifiability problems just like Poisson Background model. Specifically, *γ*_*k*_, *k* = 1, …, *K, γ*_*t*_, ***α***_0_, ***α*** and *γ*_*k*_/*c, k* = 1, …, *K, γ*_*t*_/*c, c****α***_0_, *c****α*** give the same model for any positive constant *c*. So similarly, we impose a constraint 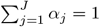 to the model.

Applying different constraints can change the parameters by a factor, while some functions of parameters, including *γ*_0_***α***_0_ and *γ*_0_***α*** are invariant to choice of constraints. Let 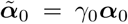 and 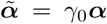, which are parameterization(constraint) invariant size factors and we use them as size factor in normalization.

### 0.3 Diagnostics and Remedy for Poisson Background model

One important metric of any Poisson model is the dispersion, i.e. the ratio of variance vs mean, which equals to 1 theoretically. In Poisson Background model, dispersion of *i*th feature and *j*th sample is

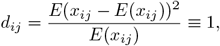

which can be estimated by squared Pearson residual at *x*_*ij*_

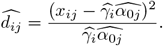

To reduce variation caused by randomness of a single point, we calculate the empirical (mean) dispersion as a diagnostics metric for this model.

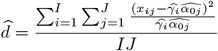

We call the data overdispersed when 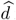 is much larger than 1, a rule of thumb cutoff can be 2. Overdispersion is a sign of extra variation not captured by model (1).

Besides empirical dispersion, a ppplot is generated as well. Let *F* (*x*|*λ*) be the cumulative distribution function of the Poisson distribution, by model assumption

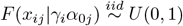

We calculate

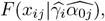

the empirical cumulative probability function values, then sort them from the smallest to the largest, and plot the sorted vector against a even grid over [0, 1] with *IJ* points, the theoretical cumulative probability function values. In practice, we have found the count data for negative probes could be very low, a decent amount could just be 0, and the ppplot would end up with zigzag shape on the lower end due to that. A way to mitigate that is to simulate data from the fitted model, generate empirical cumulative probability function values, sort them and use the sorted vector as the theoretical cumulative probability function values.

For a good model fit, the ppplot is almost a diagonal line from 0 to 1. The most common aberration of model assumption results in an “S” shape in ppplot, indicating overdispersion.

For real targets with biological variability, fitting Poisson Background model is inadequate and will lead to overdispersion. More complicated models explained in the following sections are needed to account for their additional variation.

Even the features are only negative probes, overdispersion could still occur. For data from multiple sources, mean expression level of each probe could be different. For example, if there is some dramatic difference in experimental setting among different slides of samples, the mean expression level of each probe could be different from slide to slide. Assuming there are *s* = 1, …, *S* different sources of data, a quick check is to fit the alternative model below and recalculate 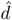 or generate heatmap using *γ*_*is*_.

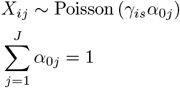

where *γ*_*is*_ are the mean expression level for *i*th feature from *s*th source (batch, slide,… etc.). A much smaller 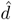 and a clear clustering in the heatmap of feature factors indicate batch effect.

Overdispersion could also occur due to outliers. We detect outliers by the empirical probability of each point using the estimated parameters. Detected outliers are mostly large outliers for negative probes. On the lower end, 0 is unlikely to be determined as outliers in a negative probe count matrix, of which counts are usually low. After outliers are identified, they can be directly set as missing and Poisson background model could be refitted with missing values. Furthermore, if *i*th feature or the *j*th ROI have a lot of counts are outliers, the *i*th feature or the *j*th sample should be considered as an outlier, and be removed from the Poisson Background model. Furthermore, the *j*th sample outlier should be removed universally from all downstream analysis since the underlying mechanism impact the negative probes could impact other features too.

### 0.4 Background Score Test

#### 0.4.1 Without prior

After applying the Poisson Background Model to the negative probes, it is imperative to test whether a target is above the background use the fitted model as a reference. Such test is called background score test.

For *k*th feature, assuming it has only one probe, and it still follows the Poisson background model

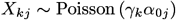

where the *α*_0*j*_ are background size factors estimated from Poisson background model on negative probes

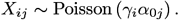

Let 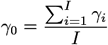, we are interested in testing

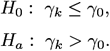

Let ***x***_*k*_ be the vector of *x*_*ik*_, the observed count for *k*th feature in *i*th sample.

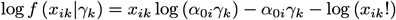

and the likelihood function

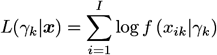

The score, i.e, the gradient of the log-likelihood function:

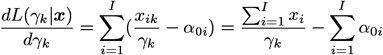

The Fisher information

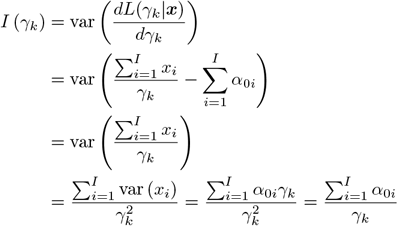

the score statistic

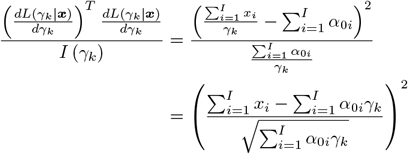

Under null hypothesis

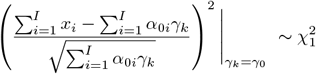

Or

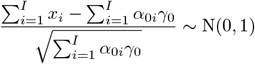

Since this is a one-sided test, we reject the null when

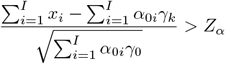

where *α* is the significance level, default is *α* = 0.001.

#### 0.4.2 With prior

For *k*th feature, assuming it has only one probe, and it still follows the Poisson background model with Gamma prior

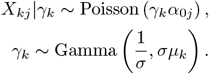

With *α*_0*j*_, *σ* and *µ*_0_ estimated from Poisson background model using negative probes

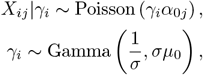

we are interested in testing

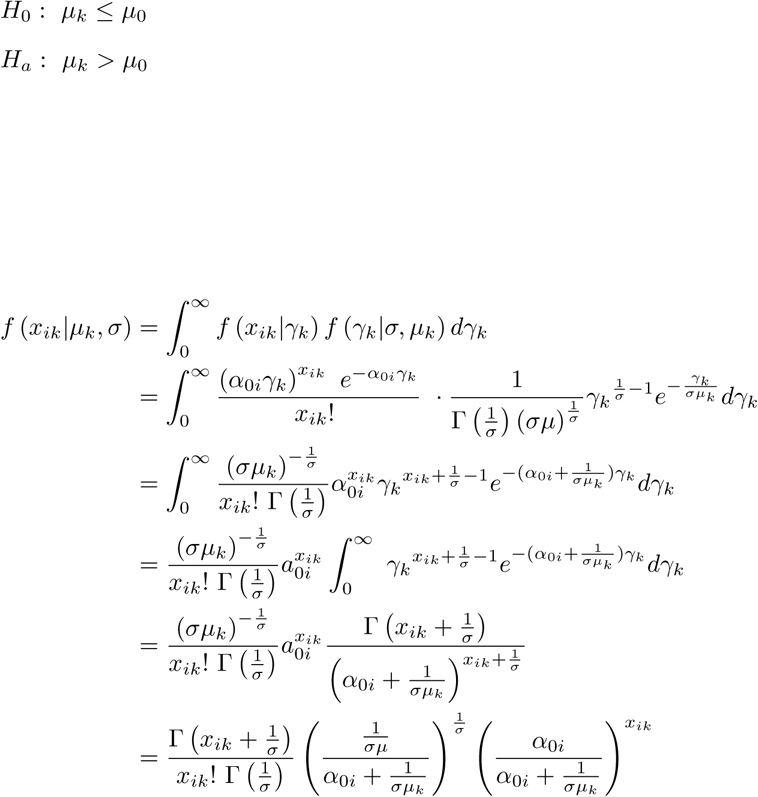

Thus

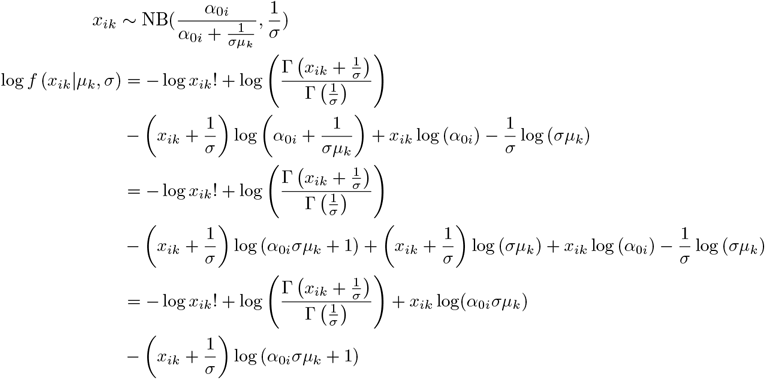

Given *σ*, the likelihood function is

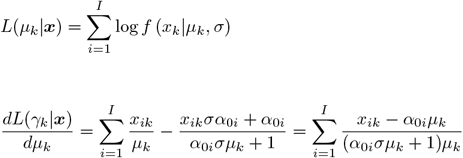

The Fisher information

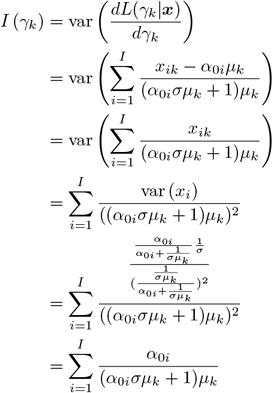

Under the null hypothesis

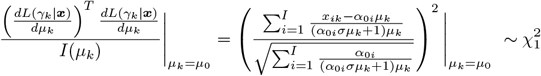

Or

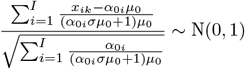

and we reject the null when

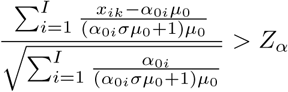

where *α* is the significance level, default is *α* = 0.001.

### 0.5 Deriving 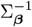 in Poisson threshold model

The covariance matrix 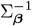 is determined by Empirical Bayes approach in 2 steps.

1. 1 Solving this model using the set of high features using a default prior 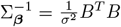, where 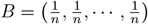. This means we are only adding a penalty of 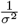 to the mean of each of these high abundance feature, the default is *σ* = 5. This prior amounts to a belief of mean log_2_ expression of each feature follows distribution 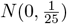. It is a weak penalty helps with the numerical stability especially with 0 counts.
2. Using the 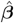 estimated from high features, calculate 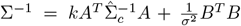, where 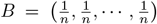, and *A*_(*n−*1)*×n*_ is any full rank matrix satisfying *AB*^*T*^ = **0**. By definition, *A*_(*n−*1)×*n*_ consists of vectors of contrasts. 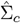 is the empirical covariance matrix of 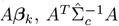 is invariant with respect to different choices of A. The contrast factor *k* ∈ (0, 1) can adjust the penality level of contrast. This form of EB prior is based on the idea of decomposing the precision matrix into the orthogonal space of precision matrix of mean and precision matrix of contrasts, rooted from the belief the contrast information of the high features should be passed to the parameter estimation of other features, but they are in different dynamic range so the behavior of mean expression of high features should not impact the behavior of other features.

It is more natural to assume only the ROIs in the same slide are correlated, so in the case of multiple slides data, it is advisable to apply this normalization function on each slide separately, which is implemented in **GeoDiff** as default for normalization of multiple slides data.

## Acknowledgements

Text for this section…

## Abbreviations

Text for this section…

## Availability of data and materials

Text for this section…

## Ethics approval and consent to participate

Text for this section…

## Consent for publication

Text for this section…

## Authors’ contributions

Text for this section …

## Authors’ information

Text for this section…

## Author details

NanoString Technologies, Seattle, US.

## Additional Files

Additional file 1 — numerics.pdf

Description of numerical methods for solving the models included in the paper

