## Supplemental Material for "Background modeling, Quality Control and Normalization for GeoMx RNA data with *GeoDiff*"

### Numerics

#### 1 Overview

All models in **GeoDiff** are solved by maximum likelihood(MLE). Only Poisson Background model has closed form MLE solution (in presence of missing values, closed form MLE solution are conditioned on other parameters fixed) and are solved by directly using these forms. Other models are solved numerically by L-BFGS-B implemented by optim function. The threshold parameter $\gamma_k$, the size factors $\boldsymbol{\alpha}$ and parameter $r$ in Negative Binomial distribution are all bounded below by 0. It is very convenient to incorporate such box contraint in L-BFGS-B.

#### 2 Poisson Background model

The log-likelihood function

$$L(\gamma_i, \alpha_j | (x_{ij})_{I\times J}) = \sum_{i=1}^{I}\sum_{j=1}^{J}(x_{ij}\log(\gamma_i\alpha_j) - \gamma_i\alpha_j - \log(x_{ij}!))$$

$$\frac{\partial L(\gamma_i, \alpha_j)}{\partial \gamma_i} = \sum_{i=1}^{I}\sum_{j=1}^{J}(\frac{x_{ij}}{\gamma_i} - \alpha_j) = 0$$

$$\frac{\partial L(\gamma_i, \alpha_j)}{\partial \alpha_j} = \sum_{i=1}^{I}\sum_{j=1}^{J}(\frac{x_{ij}}{\alpha_j} - \gamma_i) = 0$$

$$\gamma_i = \frac{\sum_{j=1}^{J} x_{ij}}{\sum_{j=1}^{J}\alpha_j}$$

$$\alpha_j = \frac{\sum_{i=1}^{I} x_{ij}}{\sum_{i=1}^{I}\gamma_i}$$

With the constraint $\sum_{j=1}^{J} \alpha_j = 1$,

$$\gamma_i = \sum_{j=1}^{J} x_{ij}$$

$$\alpha_j = \frac{\sum_{i=1}^{I} x_{ij}}{\sum_{i=1}^{I} \sum_{j=1}^{J} x_{ij}}$$

Thus the Poisson Background modeling has closed form solution with complete data. However, we often refit this model after identify a set of outliers. With a subset $\Omega$ of $(x_{ij})$ of missing values, estimation of $\gamma_i$ and $\alpha_j$ are iterative.

With the same constraint $\sum_{i=1}^{I} \alpha_i = 1$, iterate the following steps until converge

Starting value $\alpha_j = \sum_{i=1}^{I} x_{ij} I(x_{ij} \notin \Omega)$ and rescaled so that $\sum_{i=1}^{I} \alpha_i = 1$.

1. For $i = 1, \cdots, I$ calculate
$$\gamma_i = \frac{\sum_{j=1}^{J} x_{ij} I(x_{ij} \notin \Omega)}{\sum_{j=1}^{J} \alpha_j I(x_{ij} \notin \Omega)},$$
2. For $j = 1, \cdots, J$ calculate
$$\alpha_j = \frac{\sum_{i=1}^{I} x_{ij} I(x_{ij} \notin \Omega)}{\sum_{I=1}^{I} \gamma_i I(x_{ij} \notin \Omega)},$$
3. with $C = \sum_{j=1}^{J} \alpha_j$, update
$$\alpha_j = \frac{\alpha_j}{C}$$
for $j = 1, \cdots, J$.

When data is complete, thus $\Omega$ is empty, the results of this iterative algorithm are identical to closed form solution. Therefore, this algorithm is implemented to handle all scenarios.

For Poisson Background model with multiple sources, the algorithm is very similar. Let $\Phi$ be the function mapping an sample to its source, i.e. $\Phi(j) = s$ if $j$th sample is from $s$th source. Starting value $\alpha_j = \sum_{i=1}^{I} x_{ij} I(x_{ij} \notin \Omega)$ and rescaled so that $\sum_{i=1}^{I} \alpha_i = 1$.

1. For $s = 1, \cdots, S$ calculate
$$\gamma_{is} = \frac{\sum_{\Phi(j)=s} x_{ij} I(x_{ij} \notin \Omega)}{\sum_{\Phi(j)=s}^{J} \alpha_j I(x_{ij} \notin \Omega)}$$
2. For $j = 1, \cdots, J$ calculate
$$\alpha_j = \frac{\sum_{i=1}^{I} x_{ij} I(x_{ij} \notin \Omega)}{\sum_{I=1}^{I} \gamma_i I(x_{ij} \notin \Omega)}$$

3. with $C = \sum_{j=1}^{J} \alpha_j$, update

$$\alpha_j = \frac{\alpha_j}{C}$$

for $j = 1, \cdots, J$.

### 3 Negative Binomial threshold model

In the Negative Binomial threshold model

$$Y_{kj} \sim \text{NB}(\max(\gamma_k - \gamma_t, 0)\alpha_j + \max(\gamma_i, \gamma_t)\alpha_{0j}, r_k),$$

parameters $\gamma_t$, $\gamma_k, r_k, k = 1, \cdots, K$ and $\alpha_j, j = 1, \cdots, J$ are solved in the iterative fashion:

1. Optimize

$$\gamma_k, r_k, k = 1, \cdots, K$$

sequentially by fixing $\gamma_t$ and $\alpha_j, j = 1, \cdots, J$

2. Optimize

$$\alpha_j, j = 1, \cdots, J$$

sequentially by fixing $\gamma_t$ and $\gamma_k, r_k, k = 1, \cdots, K$

3. (Optional) Optimize $\gamma_t$ by fixing $\gamma_k, r_k, k = 1, \cdots, K$ and $\alpha_j, j = 1, \cdots, J$

### 4 Poisson threshold Normalization

The probability mass function for Poisson distribution with mean $\mu$ is

$$f(Y = y|\mu) = \frac{\mu^y e^{-\mu}}{y!}.$$

To make it similar to DE models, we let $X$ identity matrix and write $\boldsymbol{\beta} = X\boldsymbol{\beta}$. We use vector form to keep notation succinct. For a feature with observed count vector $\boldsymbol{y}$, the likelihood is

$$L(\boldsymbol{\beta}, \gamma|\boldsymbol{y}) = f(\boldsymbol{y}|\boldsymbol{\beta}, \gamma) = \mathbf{1}^T \left(\boldsymbol{y} \log \boldsymbol{\mu} - \boldsymbol{\mu} - \log(\boldsymbol{x}!)\right),$$

$$\text{where } \boldsymbol{\mu} = \boldsymbol{\alpha} 2^{X\boldsymbol{\beta}} + \boldsymbol{\alpha}_0 \gamma.$$

and the objective function to minimize under the priors is

$$W(\boldsymbol{\beta}, \gamma|\boldsymbol{y}, \Sigma_{\boldsymbol{\beta}}, \sigma_\gamma) = -L(\boldsymbol{\beta}, \gamma|\boldsymbol{y}) + \frac{1}{2}\boldsymbol{\beta}^T\Sigma_{\boldsymbol{\beta}}^{-1}\boldsymbol{\beta} + \frac{1}{2}\frac{(\gamma - \gamma_0)^2}{\sigma_\gamma^2}$$

$$\frac{\partial L}{\partial \boldsymbol{\mu}} = (\frac{\boldsymbol{y}}{\boldsymbol{\mu}} - \mathbf{1})^T$$
$$\frac{\partial \boldsymbol{\mu}}{\partial \boldsymbol{\beta}} = (\log 2)\mathrm{diag}(\boldsymbol{\alpha}2^{X\boldsymbol{\beta}})X$$
$$\frac{\partial \boldsymbol{\mu}}{\partial \gamma} = \boldsymbol{\alpha}_0$$

$$\begin{aligned}
\frac{\partial W}{\partial \boldsymbol{\beta}} &= -\frac{\partial L}{\partial \boldsymbol{\mu}}\frac{\partial \boldsymbol{\mu}}{\partial \boldsymbol{\beta}} + \boldsymbol{\beta}^T\Sigma_{\boldsymbol{\beta}}^{-1} \\
&= -(\log 2)\left((\frac{\boldsymbol{y}}{\boldsymbol{\mu}} - \mathbf{1})\boldsymbol{\alpha}2^{X\boldsymbol{\beta}}\right)^T X + \boldsymbol{\beta}_k^T\Sigma_{\boldsymbol{\beta}}^{-1} \\
\frac{\partial W}{\partial \gamma} &= -\frac{\partial L}{\partial \boldsymbol{\mu}}\frac{\partial \boldsymbol{\mu}}{\partial \gamma} + \frac{(\gamma - \gamma_0)}{\sigma_\gamma^2} \\
&= -(\frac{\boldsymbol{y}}{\boldsymbol{\mu}} - \mathbf{1})^T\boldsymbol{\alpha}_0 + \frac{(\gamma - \gamma_0)}{\sigma_\gamma^2}
\end{aligned}$$
